# Acute tuft cell ablation induces malabsorption and alterations in secretory and immune cell lineages in small intestine

**DOI:** 10.1101/2024.09.18.613746

**Authors:** Michael Momoh, Francisca Adeniran, Cynthia Ramos, Kathleen E. DelGiorno, Hiroshi Seno, Joseph T. Roland, Izumi Kaji

## Abstract

**Background & Aims:** Intestinal tuft cells have recently been the interest of studies in several human gastrointestinal diseases. However, the impact of tuft cell deletion on intestinal physiological functions are not fully understood. This study investigated the effects of acute tuft cell loss on nutrient absorption and cell lineage differentiation.

**Methods:** Tuft cell deletion was induced in *DCLK1-IRES-GFP-CreERT2/+;Rosa-DTA* (DCLK1-DTA) mice by a single tamoxifen injection concomitant with littermate controls. Intestinal tissues were analyzed two-, four-, or seven-days post tamoxifen injection.

**Results:** DCLK1-DTA mice showed significantly shortened small intestinal length and body weight loss on day 4. Impaired activities of Na^+^-dependent glucose transporter 1 (SGLT1) and cystic fibrosis transmembrane regulator (CFTR) were observed in Ussing chamber experiments. Tissue immunostaining revealed a transient deletion of intestinal and biliary tuft cells, which was maximal on day 4 and recovered by day 7. On day 4 post tamoxifen, cholecystokinin (CCK)+ enteroendocrine cell numbers were increased particularly in the ileum. Correlated with the tuft cell reduction, the frequency of mislocalized Paneth cells, which were co-labeled by Paneth and goblet cell markers, was increased in the villus regions. In the lamina propria, fewer mast cells and leukocytes were found in the day 4 DCLK1-DTA mice than in controls.

**Conclusion:** Ablation of intestinal tuft cells may induce nutrient malabsorption through alterations in epithelial cell proliferation and differentiation along with changes in mucosal defense response. These observations elucidate a new role for tuft cells in regulating intestinal absorption and mucosal regeneration.

## INTRODUCTION

Tuft cells are rare chemosensory cells found in many organ systems including the gastrointestinal (GI) tract. Known for their distinct bottle-shaped appearance with a ‘tuft-like’ bundle of microvilli on the apical surface, defined by the expression of the chemosensory cell-specific transcription factor POU domain, class 2, transcription factor 3 (POU2F3) (1, 2). In mouse tissues, tuft cells are marked by the neuronal protein, doublecortin-like kinase 1 (DCLK1) (3). Although this marker is shared with other cell types, DCLK1 in epithelial tissues has become an established marker for tuft cells in mice (4, 5). Tuft cells in digestive organs also express choline acetyltransferase (ChAT), which is an essential enzyme for synthesizing the neurotransmitter, acetylcholine (ACh) (6). Non-neuronal ACh release from the tuft cells is an important secretagogue for activating epithelial defense responses (7, 8). Intestinal tuft cells possess similar characteristics to taste cells utilizing key components involved in the taste signaling cascade, including transient receptor potential channel M5 (TRPM5), alpha-gustducin, and taste receptor type 1 members (TAS1Rs) (9). Whole-body knockout of alpha-gustducin or TAS1R3 in mice indicate that such sugar sensors regulate glucose transporter expression in the small intestine (10). However, the involvement of tuft cells in this regulatory mechanism has not been clarified.

In the intestinal mucosa, tuft cells are implicated in a sensory role in initiating a type 2 immune response upon challenge by a parasitic helminth or microbiota dysregulation (11, 12). Genetically modified mice lacking the chemosensory pathway (TRPM5 or alpha-gustducin deficient strains) fail to activate a type 2 immune response upon parasite colonization (13). Activated tuft cells secrete the cytokine Interleukin 25 (IL-25), which stimulates tissue resident group 2 innate lymphoid cells (ILC2s) in the lamina propria to produce IL-13. IL-13 stimulates the proliferation of the stem cells in the crypts, inducing the differentiation of more tuft cells and goblet cells (11, 13). This feed-forward circuit leads to parasite clearance in the intestinal mucosa through increased tuft cell-derived ACh and goblet cell mucin secretion. The absence of sensory tuft cells leads to a higher parasite burden in mice, significantly slowing down the “weep and sweep” process (11, 14). Yet, the interaction between intestinal tuft cells and other types of mucosal immune cells is not fully understood.

The importance of tuft cells in mucosal homeostasis is highlighted in several human gastrointestinal diseases. Indeed, tuft cell numbers are diminished in inflammatory intestinal diseases, such as ulcerative colitis and duodenitis (15, 16). Interestingly, in some cases, such diseases have been ameliorated upon tuft cell restoration as seen in a mouse model with Crohn’s-like ileitis (17). Differentiation of intestinal tuft cells may serve an important role in mucosal regeneration, while their depletion may contribute to the malabsorption seen in patients suffering from celiac disease or microvillus inclusion disease (MVID), consistent with our observations in Myosin VB knockout mice, which model MVID phenotypes (15, 18, 19). Considering the role tuft cells play in chemosensation, loss may directly induce nutrient malabsorption. To test this hypothesis, this study evaluated the changes in intestinal absorptive function, secretory cell lineages, and mucosal immune cell populations when tuft cells are acutely deleted in a tamoxifen-induced mouse model.

## MATERIALS and METHODS

### Mice

The Institutional Animal Care and Use Committee (IACUC) of Vanderbilt University Medical Center, Nashville, TN, USA approved all experimental procedures and animal care (M2000104). Frozen sperm from *DCLK1-CreER^T2-IRES-GFP^* mice (20) was a gift from Dr. Hiroshi Seno at Kyoto University and used for *in vitro* fertilization at the Vanderbilt Genome Editing Resource. After backcrossing with wild type C57Bl6/J (Jackson Laboratory), *DCLK1-CreER^T2-IRES-GFP^* mice were crossbred with ROSA26-DTA mice (Strain #009669, Jackson Laboratory). *DCLK1-CreER^T2-IRES-GFP^* and *DTA* alleles were maintained as heterozygous on C57Bl6/J to obtain *DCLK1-CreER^T2-IRES-GFP/+^;Rosa26-DTA/+* (referred to as DCLK1-DTA) and control mice (*DCLK1-CreER^T2-IRES-GFP/+^* and Rosa-DTA/+) from same litters.

At 7–9 weeks of age, both male and female DCLK1-DTA mice and littermate controls received a single dose of tamoxifen citrate (80 mg/kg) by intraperitoneal (IP) injection (day 0). Body weight changes were monitored daily. On day 2, 4, or 7, mice were euthanized, and the duodenum (0–8 cm from the pyloric ring), jejunum (8 cm following the duodenum), ileum (distal 8 cm from the ileocecal junction), and colon were collected. These intestinal segments were cut open along the mesenteric border, luminal contents were washed out, and tissues were fixed in 10% neutral buffered formalin (NBF) between filter papers overnight. Each segment was rolled along its proximal to distal axis (Swiss roll) with pieces of Parafilm® and fixed in NBF for 24 more hours. Fixed tissues were then embedded in paraffin.

Germline POU2F3 knockout mice, a generous gift from Dr. Ichiro Matsumoto (Monell Chemical Senses Center), and littermate controls were maintained on a CD1 background by the DelGiorno lab at Vanderbilt University (IACUC No. M2000077). At the age of 8–12 weeks mice were euthanized, and small intestinal tissues were harvested. Tissues were fixed as described above, and paraffin sections were generated as a comparison to DCLK1-DTA mice.

### IL-25 and IL-13 treatment

Recombinant Mouse IL-25 (IL17E) (1399-IL, R&D Systems, Minneapolis, MN) and Mouse IL13 (413-ML) proteins were reconstituted in filtered PBS and stored at –80°C until use. Following tamoxifen injection, DCLK1-DTA mice were given intraperitoneal injection of IL-25 at 0.5 µg/mouse (11) or IL-13 at 2 µg/mouse/day (21). An equal amount of PBS was injected into a vehicle control group of mice.

### Immunofluorescence Staining and Imaging

Paraffin sections were cut at 4-µm-thickness at the Vanderbilt Translational Pathology Shared Resources (TPSR). Antigen retrieval was performed using a 10 mM sodium citrate buffer containing 0.05% Tween 20 (pH 6) in a pressure cooker for 15 minutes following deparaffinization and rehydration. After cooling down, slides were rinsed in 1X PBS and blocked with Dako Protein block serum-free solution (X0909, Agilent) for 1 hour at room temperature (r/t) or overnight at 4°C. The primary antibodies listed in **Supplemental Table 1** were diluted in Dako diluent with a background reducing compound (S3022) and incubated with the pre-blocked slides for overnight at 4°C. Some primary antibodies were pre-conjugated with Alexa Fluor 555 using Zenon™ Rabbit IgG Labeling Kits (Z25305, Invitrogen). Corresponding secondary antibodies conjugated with fluorescence (Jackson Laboratory) were diluted in Dako diluent (S0809) and incubated for 1 hour at r/t. The stained slides were rinsed in 1X PBS, counterstained with Hoechst 33342, and cover slips were mounted with ProLong Gold Antifade Mountant (Thermofisher Scientific). Immunofluorescence was visualized using a Zeiss Axio Imager M2 with Apotome (Curl Zeiss) and Aperio Versa 200 (Leica).

### Immunostaining for Claudin-15

Small pieces of jejunum obtained from mice were washed in PBS to remove luminal contents and immediately frozen with OCT compound. Cryosections (15-µm-thin) were fixed in methanol for 10 minutes at –20 °C. The tissue sections were pre-incubated with PBS including normal donkey serum (10%), Triton X-100 (0.1%), and bovine serum albumin (1%) for 1 hour to block non-specific antibody interactions. Rabbit anti-CLDN15 (38-9200, ThermoFisher) antibody was diluted in the blocking solution and incubated on sections for 1 hour at r/t. After rinsing slides in PBS, the sections were incubated with Cy3-conjugated donkey anti-rabbit IgG and Hoechst 33342 diluted in PBS for 1 hour at r/t. Fluorescence was visualized using a Zeiss Axio Imager M2 with Apotome.

### Whole-mount immunofluorescence imaging in gallbladder

Mouse gallbladders were isolated from the liver, cut open to flat sheets, and fixed in NBF between filter papers. Small pieces of gallbladder were rinsed and pre-blocked in PBS containing 5% normal donkey serum and 0.3% Triton X-100. Primary antibodies were diluted in the fresh blocking buffer and incubated with tissues for 2 days at 4°C. After rinsing tissues with 0.3% Triton X-100 / PBS (TBS), secondary antibodies and Hoechst 33342 diluted in TBS were incubated for 2 hours at r/t. Fluorescence was imaged using a Nikon Ti-E microscope with an A1R laser scanning confocal system (Nikon Instruments Inc., Melville, NY).

### Measurements of tissue architecture

The length of small intestinal crypts and villi and the height of colonic mucosa were measured on whole-slide images of immunostained tissues for Ki67 and ACTG1 using Aperio ImageScope software (Leica, Wetzlar, Germany). A minimum of 10 regions were measured and averaged for each mouse.

### Digital Image Analysis

The digital analysis was conducted at the VUMC Digital Histology Shared Resource. Utilizing whole-slide fluorescence images, individual tissue segments were extracted, and machine learning was performed to generate probability maps of all intestinal tissues and antibody signals, as well as tissue folds and debris (Ilastik) (22). Mucosal architecture was divided into epithelial and non-epithelial (mesenchymal and muscle) tissue, and non-tissue area (debris and glass). These probability maps were used in combination with the original 3–5 color images to segment individual cells based on their nuclei and combined membrane marker signals using in-house coded scripts (MatLab) (23). The scrips were optimized to quantify the various populations of immunofluorescence-positive (+) cells, including CHGA+, CCK+, 5-HT+, GIP+, SOM+, LYZ+, MMP7+ TFF3+, DCLK1+, or GATA3. These marker+ cells were identified and filtered by their size, location, and presence of a detectable nucleus. The percent of each immunopositive cell population was calculated per total number of epithelial cells in whole Swiss rolls.

### Ussing chamber experiments

Mucosal-submucosal preparations were obtained from the jejunum and mounted in sliders with an aperture = 0.3 cm^2^ (Physiologic Instruments, Leno, NV) as described previously (24). Gallbladders were cut-opened and mounted in sliders with an aperture = 0.031 cm^2^. Luminal and serosal surfaces of tissue were bathed in 4 ml Krebs-Ringer solution (117 mM NaCl, 4.7 mM KCl, 1.2 mM MgCl_2_, 2.5 mM CaCl_2_, 1.2 mM NaH_2_PO_4_, 25 mM NaHCO_3_, 11 Mm glucose) and maintained at 37°C using a water-recirculating heating system. Indomethacin (10 µM) was added into the serosal bath for jejunum to suppress prostaglandin-stimulated secretion at the baseline. The solution was continuously bubbled with a gas mixture of 95% O_2_ and 5% CO_2_ to maintain the pH at 7.4. Short-circuit current (*I*_sc_) was continuously recorded under voltage clamp conditions at zero potential difference by the DataQ system (Physiologic Instruments).

Transmucosal resistance (*R*t) was determined by changing the clamped voltage automatically at ±3 mV for 20 ms every 2 sec. An increase of *I*_sc_ indicates luminal-to-serosal current flow, e.g., anion secretion or cation absorption. The tissues were stabilized for 20 min before the effects of drugs were investigated. SGLT1 activity was represented by phlorizin (0.1 mM)-sensitive *I*_sc_. Cl^−^ secretion was measured by *I*_sc_ peaks after carbachol (10 µM) and forskolin (10 µM). CFTR activity was determined using the CFTR-inhibitor, (R)-BPO-27 (10 µM). DMSO < 0.3% in the bathing solution did not affect the *I*_sc_ or *R*_t_.

### RNA extraction and quantitative PCR

Total RNA was prepared from scraped mucosa of jejunum by using TRIzol (Thermo Fisher Scientific). After reverse transcription of the RNA using SuperScript III First-strand Synthesis System for RT-PCR (Thermo Fisher Scientific), cDNA was used as a template for PCR. Real-time quantitative PCR was performed utilizing SsoAdvanced Universal SYBR Green Supermix with a CFX96 Real-Time System (Bio-Rad Laboratories). Previously published primer pairs used were listed in **Supplemental Table 2**. Target gene expression was calculated as relative values by the ΔΔCt method.

### Chemicals

Tamoxifen citrate (USP, Spectrum™ Chemical 18-607-202) and 10x PBS were purchased from Thermo Fisher Scientific (Hanover Park, IL), and other chemicals were from Sigma Aldrich, St. Louis, MO unless specified. Tamoxifen citrate (20 mg/ml) was dissolved in 10% DMSO and 90% corn oil and warmed in a water bath at 65°C. Indomethacin (Cat# I7378) was dissolved in 100% ethanol. Forskolin was purchased from Cayman Chemical (Cat# 11018, Ann Arbor, MI), (R)-BPO-27 (Cat# HY-19778) and CaCCinh A-01 (Cat# HY-100611) were from MedChemExpress (Monmouth Junction, NJ) and were dissolved in DMSO to prepare 1000x stocks.

### Statistics

Statistical differences were determined using GraphPad Prism 8 with a significant *P* value of ≤ 0.05. The test used in each analysis is described in the figure legends.

## RESULTS

### Diphtheria toxin-induced acute tuft cell loss decreased nutrient absorption in the small intestine

The impact of acute deletion of tuft cells on intestinal absorptive function has not been fully explored. We investigated the effects of a transient DCLK1+ tuft cell loss on epithelial cell differentiation and nutrient absorptive ability in the small intestine. After a single dose of tamoxifen, DCLK1-DTA mice showed a decrease in body weight within four days and recovered to initial body weight by day 7 (**Figure 1A**). The body weight loss on day 4 was significant as compared to tamoxifen-treated control littermates. Consistent with the body weight loss, the small intestinal length, but not colonic length, was significantly shorter in DCLK1-DTA mice as compared with those of controls at day 4 post tamoxifen (**Figure 1B, C**). The shortening of intestinal length was restored upon administration of IL-25 to DCLK1-DTA mice (**Figure 1B).** Immunostaining for DCLK1 in the small intestines revealed that intestinal tuft cell numbers were decreased in DCLK1-DTA mice by 85% on day 2, largely abolished on day 4, and completely recovered on day 7, consistent with their body weight changes (**Figure 1D, E**). The correlation between tuft cell numbers and small intestinal length suggests that acute deletion of tuft cells may influence mucosal cell proliferation, therefore we explored this as a possibility in our mouse model.

**Figure 1.**
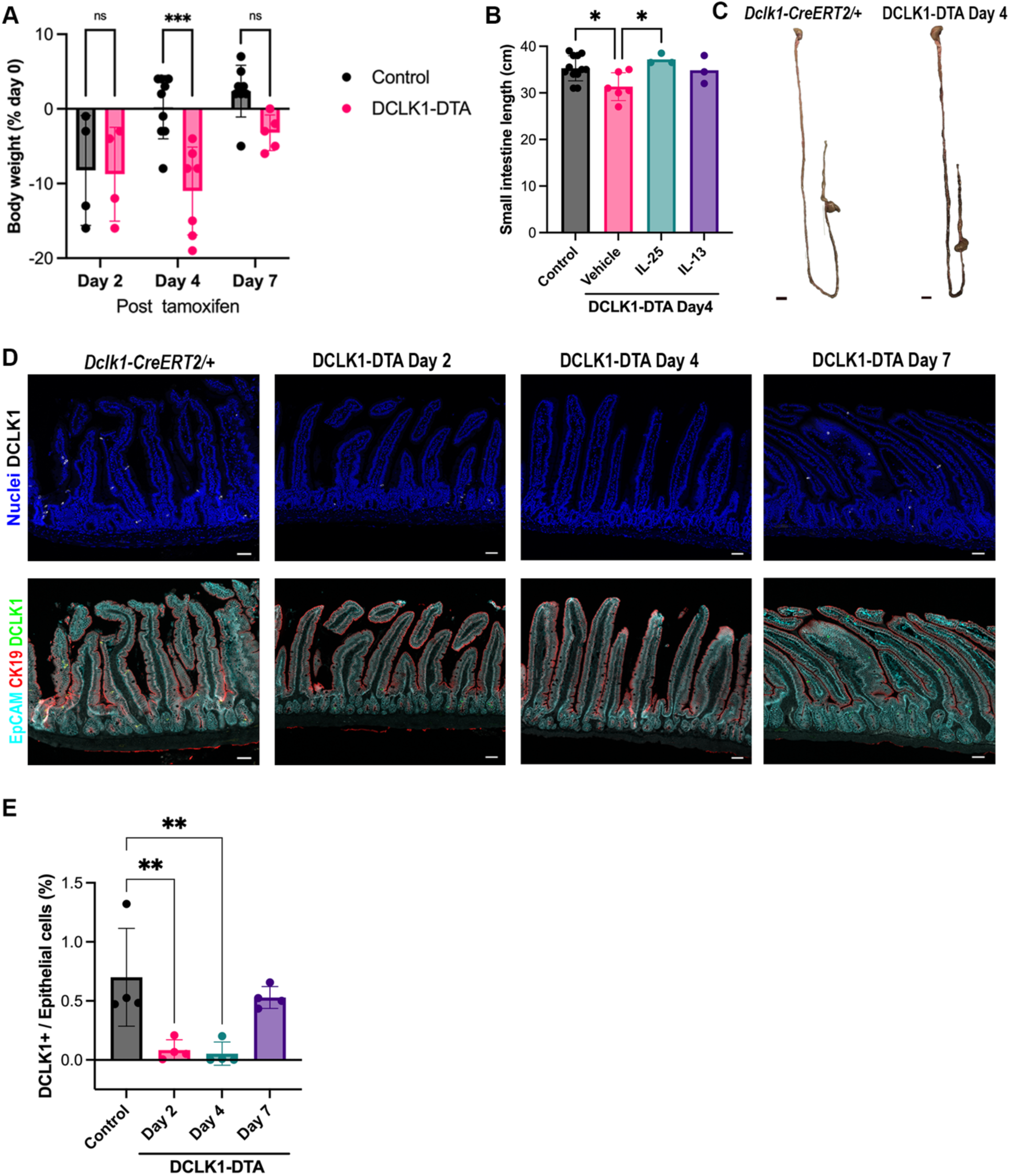
Tuft cell deletion exerts nutrient absorption deficits in mouse small intestine. **(A)** Significant body weight loss is observed only on day 4 post tamoxifen treatment in DCLK1-DTA mice versus control mice. \*\*\**P* < 0.001 by two-way ANOVA following Tukey’s multiple comparisons test. (**B)** The small intestine is significantly shortened on day 4 in DCLK1-DTA mice as compared to control mice and is restored upon IL-25 supplementation. \**P* < 0.05 by ANOVA with Dunnett’s test. **(C)** Representative images of whole GI tract from control mice and DCLK1-DTA day 4 mice. Scale bars = 1 cm. **(D)** Immunostaining for DCLK1 in jejunum sections. Epithelial markers, EpCAM and CK19, are co-stained to conduct cell counting by imaging analysis scripts. Compared to healthy control mice, DCLK1-DTA mice demonstrate transient decreases in tuft cells after a tamoxifen injection. Scale bars = 50 µm. **(E)** Quantification of DCLK1+ cells per total epithelial cell numbers. More than 95% deletion occurs in DCLK1-DTA mice on day 4. \*\**P* < 0.01 ANOVA with Dunnett’s test. In all graphs, each datapoint indicates individual mouse, and bars show Mean ± S.D..

Immunostaining for a proliferation marker, Ki67, revealed a decrease in the length of Ki67+ crypts in DCLK1-DTA mice on day 4, whereas villus length was not significantly affected between DCLK1-DTA and control tissues (**Figure 2A, B**). This significant decrease in Ki67+ crypt length was also identified in the proximal colon, but not in the distal colon (**Figure 2C, D**). To evaluate whether the effect of tuft cell loss on proliferative cells is consistent in a constitutive mouse model of tuft cell ablation, we immunostained for Ki67 in whole-body *Pou2f3* knockout mice (POU2F3^―/―^), which lack the Pou domain class 2 transcription factor 3, the master regulator transcription factor for tuft cell development. Ki67+ crypt and villus lengths were comparable between POU2F3^―/―^ and wild-type (WT) littermates (**Figure 2E, F**). Body weight of POU2F3^―/―^ mice and age-matched littermate controls had no significant difference (data not shown). These observations indicate that acute tuft cell deletion affects mucosal homeostasis and nutrient absorption of the small intestine and proximal colon, while germline POU2F3 ablation on outbred mouse background might possess a compensation pathway to maintain epithelial proliferation. The DCLK1-DTA mouse tissues on day 4 (referred to as DCLK1-DTA) were then analyzed in subsequent experiments.

**Figure 2.**
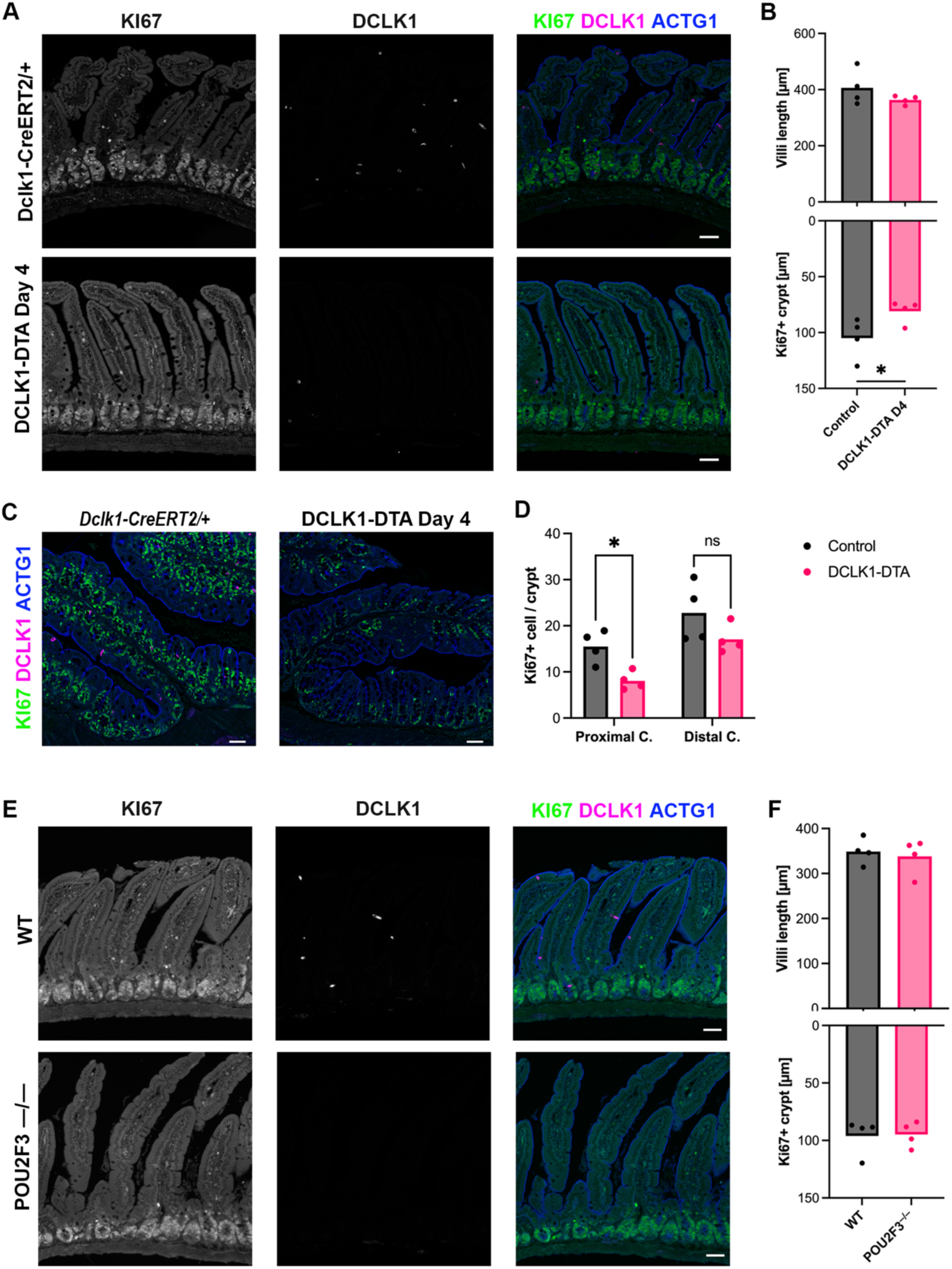
Reduced intestinal proliferation in tuft cell deleted mice. (**A and C**) Immunostaining for a proliferative cell marker, Ki67 (green), an epithelial-dominate actin, gamma-actin (ACTG1, blue), and DCLK1 (magenta) in the duodenum (A) and proximal colon (C) of DCLK1-DTA mice on day 4. **(B and F)** Ki67+ crypt length and villus height are measured in 10 regions of jejunal Swiss roll of each mouse. Each datapoint indicates an average value of individual mouse, and the bar graphs represent mean values of group. Decreased crypt proliferation is characterized in DCLK1-DTA mice, but not POU2F3–/– mice, compared to littermate control mice. \**P* < 0.05 by unpaired *t*-test. **(D)** Ki67+ nuclei are counted per vertical section of crypt in proximal and distal colons. Significant reduction of proliferative cells is identified in the proximal but not distal colon of DCLK1-DTA mice. \**P* < 0.05 by two-way ANOVA with LSD test. **(E)** POU2F3^―/―^ mice show no significant difference in Ki67+ crypt length or villus structures when compared to control littermates. Scale bars = 50 µm.

We investigated whether the absorptive function of the intestine was affected by acute tuft cell deletion. Epithelial transporter function was assessed for the sodium-dependent glucose transporter 1 (SGLT1), cystic fibrosis transmembrane regulator (CFTR), and calcium Activated Chloride Channel (CaCC) in mouse jejunum utilizing Ussing chambers. Baseline short circuit current (*I*_sc_) in steady state was significantly lower in DCLK1-DTA mice as compared to control mice jejunum (**Figure 3A**). In contrast, transmural mucosal resistance (Rt) was comparable at steady state (**Figure 3B**). The contributions of SGLT1, CFTR, and CaCCs to the baseline *I*_sc_ were evaluated by using selective inhibitors against each transporter. SGLT1-mediated absorptive current was induced by 11 mM glucose in the luminal buffer and was inhibited by phlorizin. This absorptive current was significantly reduced in DCLK1-DTA mice by approximately 80% as compared to control mice (**Figure 3C**). Chloride secretory responses were assessed using carbachol (CCh) and forskolin, which increase intracellular calcium and cAMP, respectively. While CCh-induced transient chloride secretion was comparable between DCLK1-DTA and control mice jejunum, forskolin-stimulated secretory current was decreased by 10% in DCLK1-DTA mice as compared to control (**Figure 3D**). Subsequently, CFTR and CaCC dependence in the secretory state was evaluated by adding a CFTR inhibitor, R-BPO-27, followed by a CaCC inhibitor, CaCCinh A-01. The secretory state *I*_sc_ was approximately 80% dependent on CFTR in both control and DCLK1-DTA mice, which was significantly larger than the CaCC-dependent portion (**Figure 3E**). The observed decreases in sodium absorption and chloride secretion suggest a significant change in nutrient absorption ability in the small intestine following tuft cell depletion.

**Figure 3.**
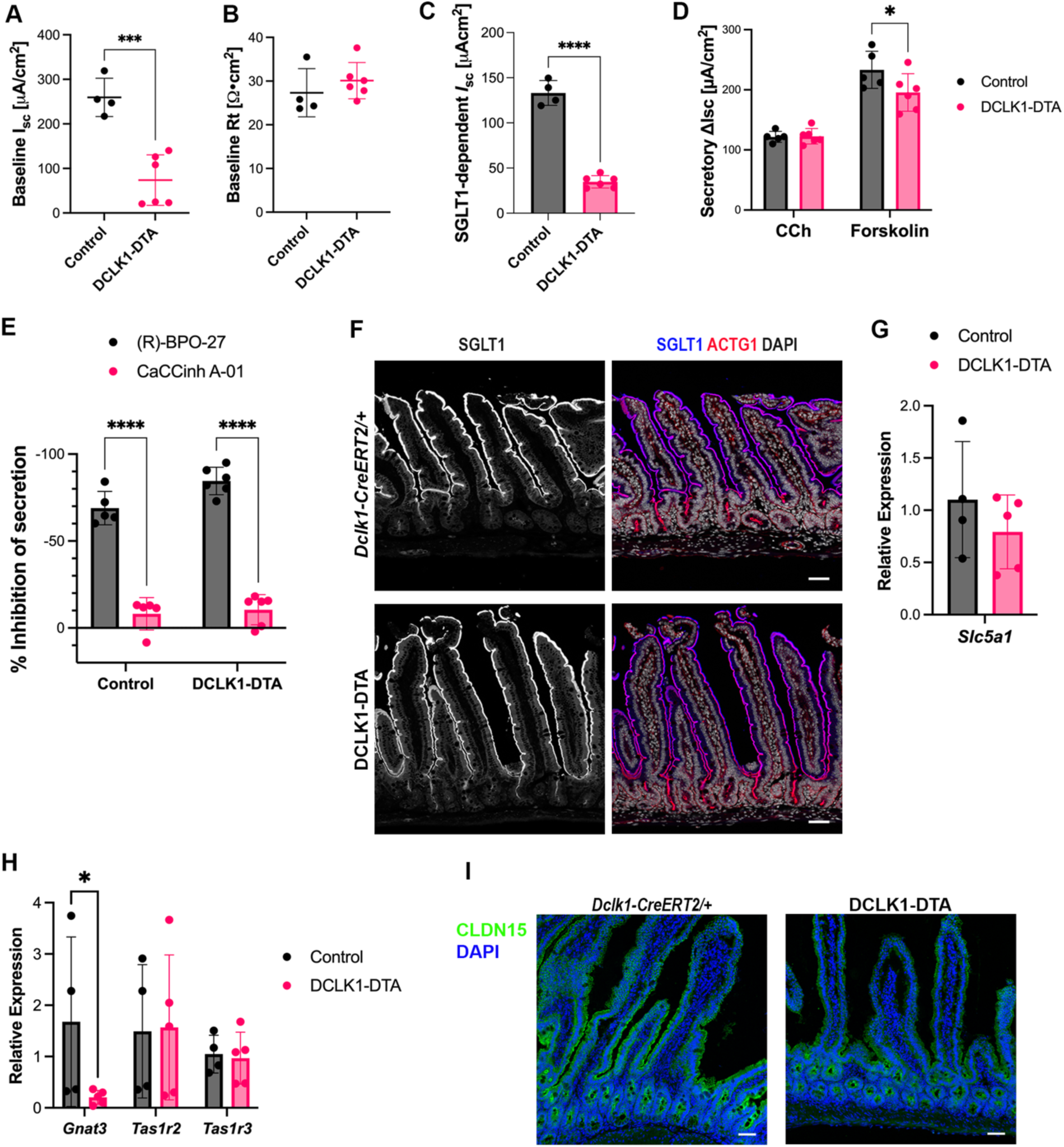
SGLT1 and CFTR functions in DCLK1-DTA mouse jejunum. (**A–E**) Epithelial ion transport measurements in jejunum utilizing Ussing chambers. Baseline short-circuit current (*I*_sc_) (A) is significantly lower in DCLK1-DTA day 4 mice than in that of control, whereas baseline transmural resistance (R_t_) (B) is comparable. DCLK1-DTA mice demonstrate significantly lower levels of SGLT1-mediated absorption (C) and forskolin-stimulated, but not carbachol (CCh)-mediated secretion (D), compared to control. Dependency on CFTR and CaCC of the stimulated secretion (E) have no statistical difference between genotypes. \**P* < 0.05, \*\**P* < 0.01, \*\*\**P* < 0.001 by unpaired t-test in graphs A–C, by two-way ANOVA with Tukey’s test in graphs D and E. **(F)** Immunostaining for SGLT1 (blue) and ACTG1 (red) in jejunum. SGLT1 is colocalized with ACTG1 on the apical brush border of villus epithelial cells in DCLK1-DTA mice and control. **(G)** Relative expression of mRNA calculated with ΔΔCt method. No significant difference is detected by Mann-Whitney test in *Slc5a1* (SGLT1) expression of jejunum mucosa between DCLK1-DTA mice compared to control. Each datapoint indicates an individual mouse. **(H)** Relative expression levels of glucose sensors. *Gnat3* (Gustducin) is significantly reduced in DCLK1-DTA day 4 mice compared to control \**P* < 0.05 by two-way ANOVA with LSD test. **(I)** Immunostaining for Claudin-15 (CLDN15, green) in jejunum. CLDN15 is localized on the tight junctions of epithelial cells and no discernible difference was identified between DCLK1-DTA mice and control. Scale bars = 50 µm.

The molecular machinery for glucose absorption was evaluated, including SGLT1, alpha-gustducin, and CLDN15. To assess the localization of SGLT1, we immunostained murine small intestinal sections for SGLT1. SGLT1 consistently colocalized with the brush border marker, gamma-actin (ACTG1), in villus epithelial cells in both DCLK1-DTA and control mice (**Figure 3F**). CLDN15 is a cation channel-forming claudin protein, involved in SGLT1 activity through sodium recycling (25). Immunostaining for CLDN15 identified expression on the basolateral membrane of enterocytes of villi, in a pattern that is indistinguishable between DCLK1-DTA and control mouse jejunum (**Figure 3F**). These immunostaining patterns indicate that acute tuft cell deletion had no effect on localization of apical SGLT1 or basolateral CLDN15, which are important for glucose absorption. Additionally, qPCR was performed with mRNA extracted from scraped jejunal mucosa of mice for *Slc5a1*, the gene that encodes SGLT1, *Gnat3*, which encodes alpha-gustducin, and two sweet taste receptors, *Tas1r2* and *Tas1r3*. The transcription level of *Slc5a1* was comparable between DCLK1-DTA and control mice (**Figure 3G**). *Gnat3* expression was significantly reduced in the jejunal mucosa of DCLK1-DTA mice as compared to control, whereas *Tas1r* members were similar to control (**Figure 3H**). Gustducin is a chemosensory molecule in taste buds shown to also be important for the sensory function of tuft cells in the small intestine (13, 26, 27). The observed reduction of *Gnat3* expression as a glucose sensor and reduced absorptive function of SGLT1 most likely corresponds to the acute deletion of tuft cells in DCLK1-DTA mice.

### Aberrant secretory cell differentiation associated with acute tuft cell loss in small intestine

Recent evidence has shown that deletion of Sox9, an integral transcription factor for Paneth cell development, increased tuft cell frequency and activity in the small intestine (28). Therefore, we investigated our DCLK1-DTA model for a connection between secretory cell lineage differentiation and tuft cells. First, we performed immunostaining for lysozyme (LYZ), an established Paneth cell marker. In addition to typical Paneth cell positioning at the base of crypts, we found LYZ+ cells further migrated toward villus tips more frequently in DCLK1-DTA mice than in control tissues (**Figure 4A**). Mislocated LYZ+ epithelial cells in villus regions were counted and compared between control and DCLK1-DTA mice for all timepoints. LYZ+ cells were rarely present in the villi of control and DCLK1-DTA murine intestines at day 2 and day 7. However, the frequency of mislocalized cells increased three to four-fold in DCLK1-DTA mice on day 4, correlating with total tuft cell deletion (**Figure 4B**). Periodic Acid-Schiff (PAS)-staining on the same sections illustrated that these cells have a goblet cell-like morphology: diffuse cellular staining as opposed to the tighter granular pattern in typical Paneth cells (**Figure 4A**). This result led us to hypothesize that this cell population may possess mixed characteristics of Paneth and goblet cells. Interestingly, mislocalized LYZ+ secretory cells were also present in the POU2F3^―/―^ mice, indicating that the formation of these cells was most likely connected to the absence of tuft cells (**Figure 4C**).

**Figure 4.**
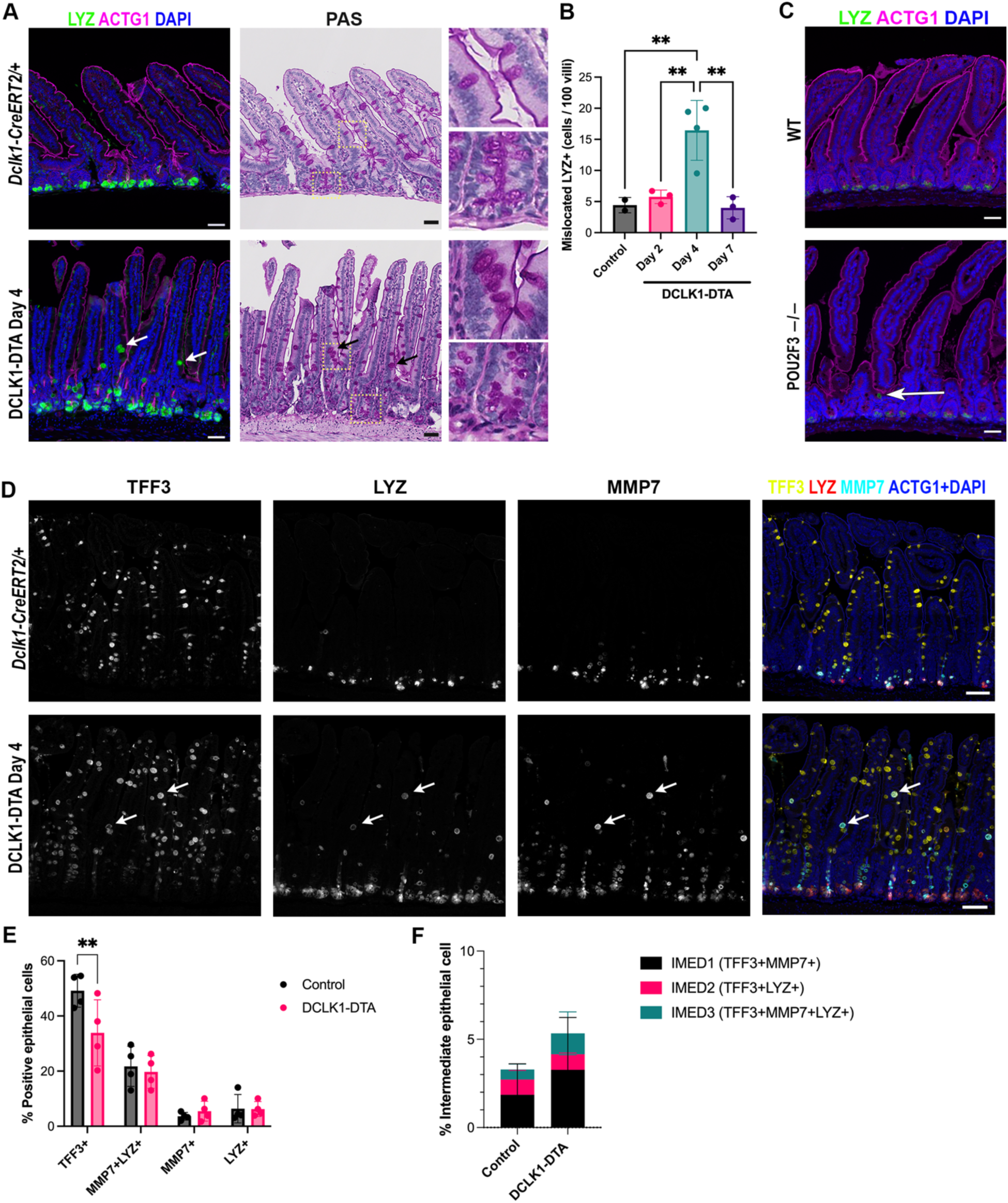
Mislocalized Paneth cells are present in acute tuft cell deletion and germline tuft cell loss models. **(A)** Immunostaining for LYZ (green) and ACTG1 (magenta) in jejunum. LYZ+ Paneth cells are limited to the bottom of crypts in control tissues, whereas DCLK1-DTA mice possess LYZ+ cells in upper crypt and villus regions (arrows). PAS staining was performed on the same slides after immunostaining. Insets show LYZ^−^ goblet cells and LYZ+ Paneth cells in control and LYZ+ mislocalized cells in DCLK1-DTA mice. Scale bars = 50 µm. **(B)** LYZ+ cells localized out of the crypt bottom are counted among 100 villi of jejunum. These cells appear more frequently in DCLK1-DTA mice at day 4 post tamoxifen compared to day 2, day 7, and control. \*\**P* < 0.01 by ANOVA with Tukey’s test. Each datapoint indicates an individual mouse. **(C)** POU2F3^―/―^ mouse tissues demonstrate mislocalized LYZ+ (green) cells out of crypts (arrow). **(D)** Co-immunostaining for TFF3 (yellow), LYZ (magenta), and Zenon-labeled MMP7 (cyan) in jejunum. ACTG1 and nuclei are shown in blue for visualizing tissue morphology. Triple-positive cells for all markers (white arrows) are localized in villus region of DCLK1-DTA day 4 tissues. **(E-F)** Digital image analysis for immunostained cell quantification in entire small intestine. TFF3-single positive cell number per all epithelial cells is significantly reduced in DCLK1-DTA mouse small intestine (E). There was no difference in frequency of Paneth cell marker-positive, TFF3^−^ cells. Double-or triple-positive cell numbers for both goblet and Paneth cell markers are shown as intermediate (IMED) type of cells (F). Bar graphs represent Mean ± S.D.. \*\**P* < 0.01 by two-way ANOVA with LSD test.

Therefore, we co-immunostained for a goblet cell marker, Trefoil Factor 3 (TFF3), and Paneth cell markers, matrix metalloprotease 7 (MMP7) and LYZ, as well as ACTG1 to quantify epithelial cell numbers expressing each cell marker (**Figure 4D**). In control tissues, LYZ and MMP7 staining was limited to crypt regions with intense signals in the base of the glands, while TFF3 was identified in goblet cells throughout the crypt-villus axis with intense signals in the villus regions. In the DCLK1-DTA tissues on day 4, LYZ and MMP7 expression was more frequently identified in villi than control and predominantly colocalized with TFF3. Further, digital image analysis on whole small intestinal sections identified all epithelial cells and calculated the immunostaining for each marker by combining machine learning and cell segmentation tools. The cell population of TFF3+ / LYZ– / MMP7– was significantly smaller in DCLK1-DTA tissues than in control, indicating that mature goblet cell numbers were decreased following tuft cell deletion (**Figure 4E**). There were similar numbers of TFF3-negative Paneth cell subpopulations, including double positive LYZ+ / MMP7+, and either LYZ+ or MMP7+ single-positive cells in the entire crypt-villus axis (**Figure 4E**). Cells double or triple positive for goblet and Paneth cell markers were also quantified as intermediate cell types (IMED), such as TFF3+ / LYZ+ / MMP7–, TFF3+ / LYZ– / MMP7+, and TFF3+ / LYZ+ / MMP7+ (**Figure 4F**). The observed numbers of IMED cells showed a higher trend in DCLK1-DTA mice as compared to control tissues, however, there was no statistical difference. These analyses suggest that the rare intermediate, or immature, secretory cells that express both goblet and Paneth cell markers are present in healthy control tissues, typically in the crypts. As DCLK1-DTA tissues demonstrate goblet / Paneth intermediate cells more frequently in the villi, tuft cell deletion may interrupt terminal differentiation of goblet cells.

### Tuft cell deletion reciprocally increased enteroendocrine cells in the small intestine

To test the hypothesis that these mixed-labeled secretory cells are present due to stalled differentiation in all secretory cell lineages, we investigated enteroendocrine cells (EECs). After acute tuft cell depletion, immunostaining was conducted for a pan-EEC marker, chromogranin A (CHGA) (**Figure 5A**). The CHGA+ EEC population in epithelial cell was determined by digital image analysis tools in entire small intestine and revealed a significant increase in DCLK1-DTA mice as compared to control mouse tissues (**Figure 5B**). We next sought to identify the subpopulation of increased EECs following tuft cell loss. Cholecystokinin (CCK) has been reported to play a role in helminth clearance (29). Immunostaining for CCK and digital image analysis were performed to quantify all CCK+ cells in the entire small intestine (**Figure 5C**). The results revealed that the CCK+ EEC subpopulation was significantly higher in the ileum of DCLK1-DTA mice compared to control mouse intestine and other segments of DCLK1-DTA mice (**Figure 5D**). Other major EEC subtypes were further evaluated, including serotonin+ (5-HT) enterochromaffin cells, gastric inhibitory peptide+ (GIP), and somatostatin+ (SST) (**Figure 5E**). Positive cells were digitally quantified and comparable numbers of each subpopulation were detected between DCLK1-DTA and control mice (**Figure 5F**). Taken together, these observations suggest a specialized epithelial cell remodeling pattern following acute tuft cell depletion in mouse small intestine.

**Figure 5.**
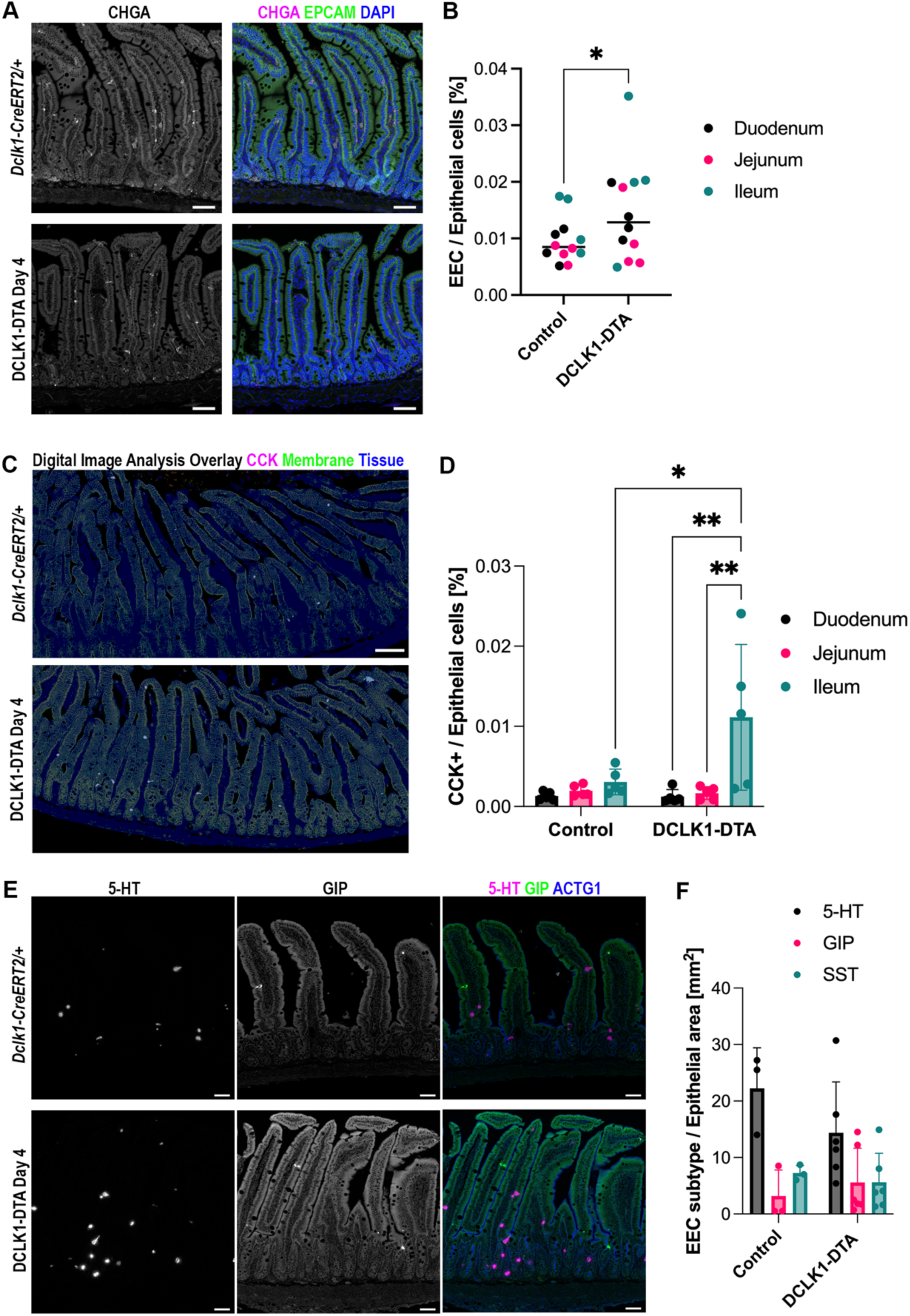
Enteroendocrine cell numbers are increased in mouse small intestine following acute tuft cell deletion. **(A)** Immunostaining for a pan-enteroendocrine cell marker, CHGA, and an epithelial cell marker, EpCAM, in small intestines of control and DCLK1-DTA mice. Scale bars = 50 µm. **(B)** EpCAM+/CHGA+ cell (EEC) numbers per all epithelial cells are significantly increased by acute tuft cell deletion. Each datapoint indicates an individual intestinal roll, and bars indicate medium. \**P* < 0.05 by unpaired *t*-test. *N* = 4 mice. **(C)** Overlay of immunostaining images for CCK and EpCAM with epithelial cell segmentation generated by digital image analysis. Light gray subjects indicate CCK+ (magenta) EECs in the tissue area (blue). **(D)** CCK+ cell frequency was significantly higher in the ileum of DCLK1-DTA cells. \**P* < 0.05, \*\**P* < 0.001 by two-way ANOVA with Tukey’s test. Each datapoint indicates an individual intestinal roll, and bar graphs represent Mean ± S.D.. *N* = 4 mice. **(E)** Co-immunostaining for EEC subtypes, 5-HT (magenta), Zenon-labeled GIP (green), an SOM (not in the fields shown), with an epithelial structural marker, ACTG1 (blue). Scale bars = 50 µm. **(F)** Positive cell numbers for 5-HT, GIP, or somatostatin (SST) per total epithelial cells are quantified by digital image analysis scripts. Each datapoint indicates an average value of entire small intestine in individual mouse. Bar graphs represent Mean ± S.D.. *N* = 3–5 mice. No statistical difference is detected between genotypes by two-way ANOVA.

### Acute tuft cell deletion alters mucosal immune cell populations in the small intestine

Next, we investigated whether changes occur in mucosal immune cell populations in the intestine following acute tuft cell deletion. We analyzed GATA binding protein 3 (GATA3), a marker of ILC2 populations, which have been shown to respond to tuft cell activity (30). Immunostaining for GATA3 was sporadically identified in the lamina propria both in crypt and villus regions (**Figure 6A**). There was no significant difference in the number of GATA3+ cells in whole Swiss rolls of small intestine between genotypes (**Figure 6B**). Immunohistochemistry was performed for CD11b, which is encoded by *Itgam* and a marker of myeloid cells, including leukocytes and dendritic cells (**Figure 6C**). Subsequent analyses utilizing QuPath (31) revealed that CD11b+ area was significantly less in the jejunum of DCLK1-DTA mice as compared to controls (**Figure 6D**). The number of mucosal mast cells increased in response to type 2 immune activation and tuft cell-derived IL-25 (32) as shown by immunostaining for mast cell protease 1 (MCPT1). MCPT1+ populations were rare in both control and DCLK1-DTA murine intestines and were localized in the lamina propria adjacent to the epithelial layer (**Figure 6E**). Their numbers were compared per 10 mm of longitudinal length of jejunum between genotypes. We identified a 50% reduction in MCPT1+ mast cell counts in the DCLK1-DTA mouse jejunum as compared to control (**Figure 6F**). We immunostained for CD3 to assess the major intraepithelial lymphocyte population, which is important for maintaining the mucosal barrier (**Figure 6E**). There was no significant change in localization or numbers of CD3+ lymphocytes following tuft cell deletion (**Figure 6G**).

**Figure 6.**
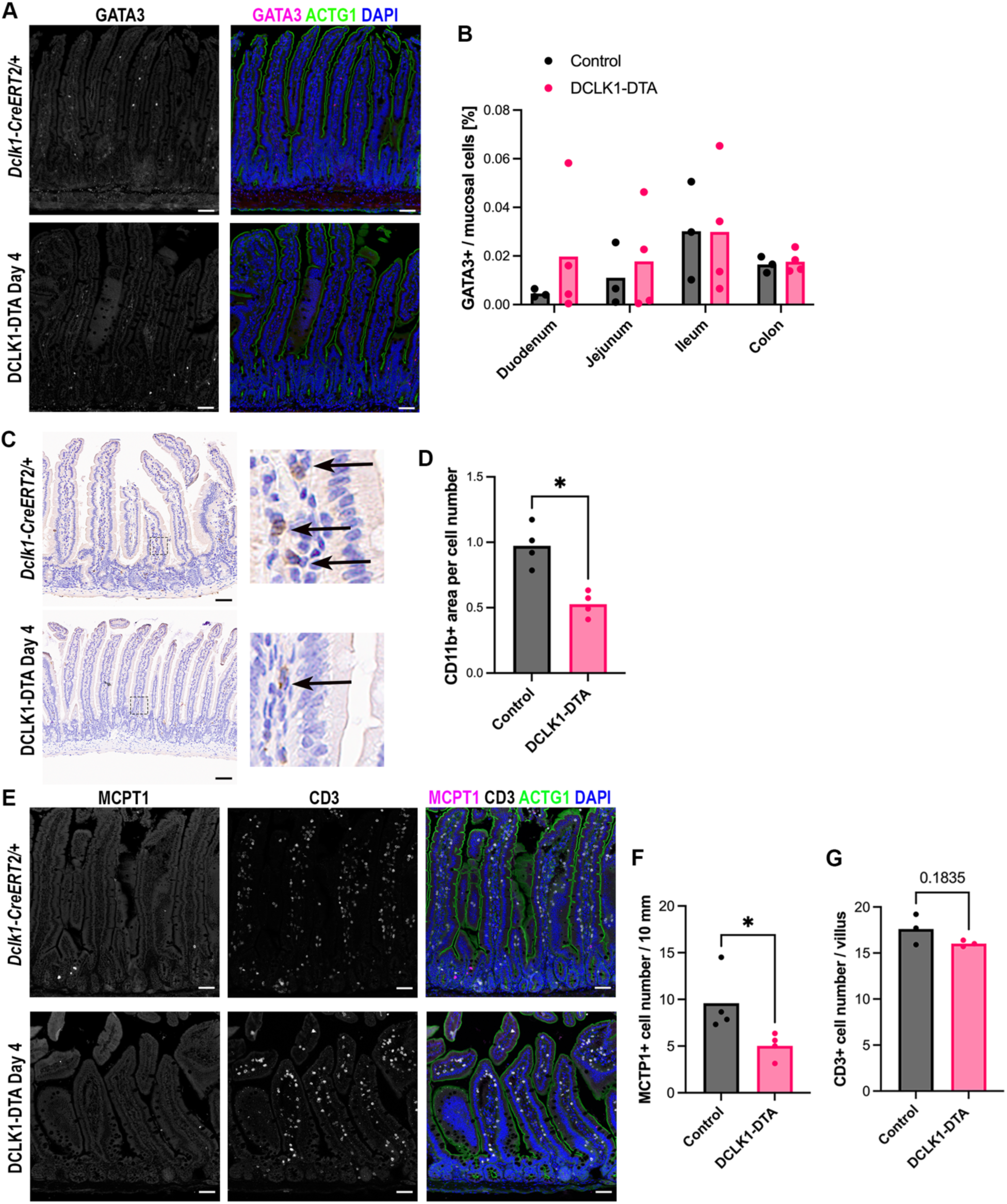
Tuft cell deletion alters mucosal immune cell populations. **(A)** Immunostaining for GATA3 (magenta), a marker for ILC2s, and ACTG1 in duodenum. GATA3 is expressed in a rare population of lamina propria cells in control and DCLK1-DTA tissues. Scale bars = 50 µm. **(B)** Digital quantification of GATA3+ cells per total mucosal nuclei in each Swiss roll of intestine. Each datapoint indicates an individual mouse. No statistical difference is detected by two-way ANOVA. **(C)** Immunohistochemistry for leukocyte marker, CD11b in duodenum with hematoxylin counterstaining for nuclei. Insets show CD11b+ immune cells in lamina propria (black arrows). **(D)** Quantification of CD11b+ cell area per mucosal cell nucleus number. DCLK1-DTA mouse tissues demonstrated significantly less CD11b+ cells than controls. Each datapoint indicates an average value of >10 images of an individual mouse. \**P* < 0.05 by Mann-Whitney test. **(E)** Immunostaining for mast cell marker, MCPT1 (magenta), a T-cell marker, CD3, and ACTG1 in duodenum. **(F)** MCPT1+ cell numbers are significantly decreased in DCLK1-DTA mouse duodenum compared to controls. \**P* < 0.05 by unpaired t-test. **(G)** CD3+ intraepithelial lymphocytes are comparable between mouse groups.

### Absence of inflammation and unchanged growth factors in shortened small intestine following tuft cell deletion

Small intestinal shortening is frequently coincident with mucosal inflammation (33). However, there was no evidence of significant inflammation (based on lymphocyte infiltration) on H&E-stained tissues from DCLK1-DTA mice. To confirm if tuft cell ablation has no impact on intestinal inflammation, we performed immunohistochemistry for macrophage/dendritic cell marker, F4/80 (also known as Adhesion G protein-coupled receptor E1: ADGRE1). No abnormal infiltration of F4/80+ immune cells was identified in DCLK1-DTA tissues as compared to controls (**Figure 7A-B**).

**Figure 7.**
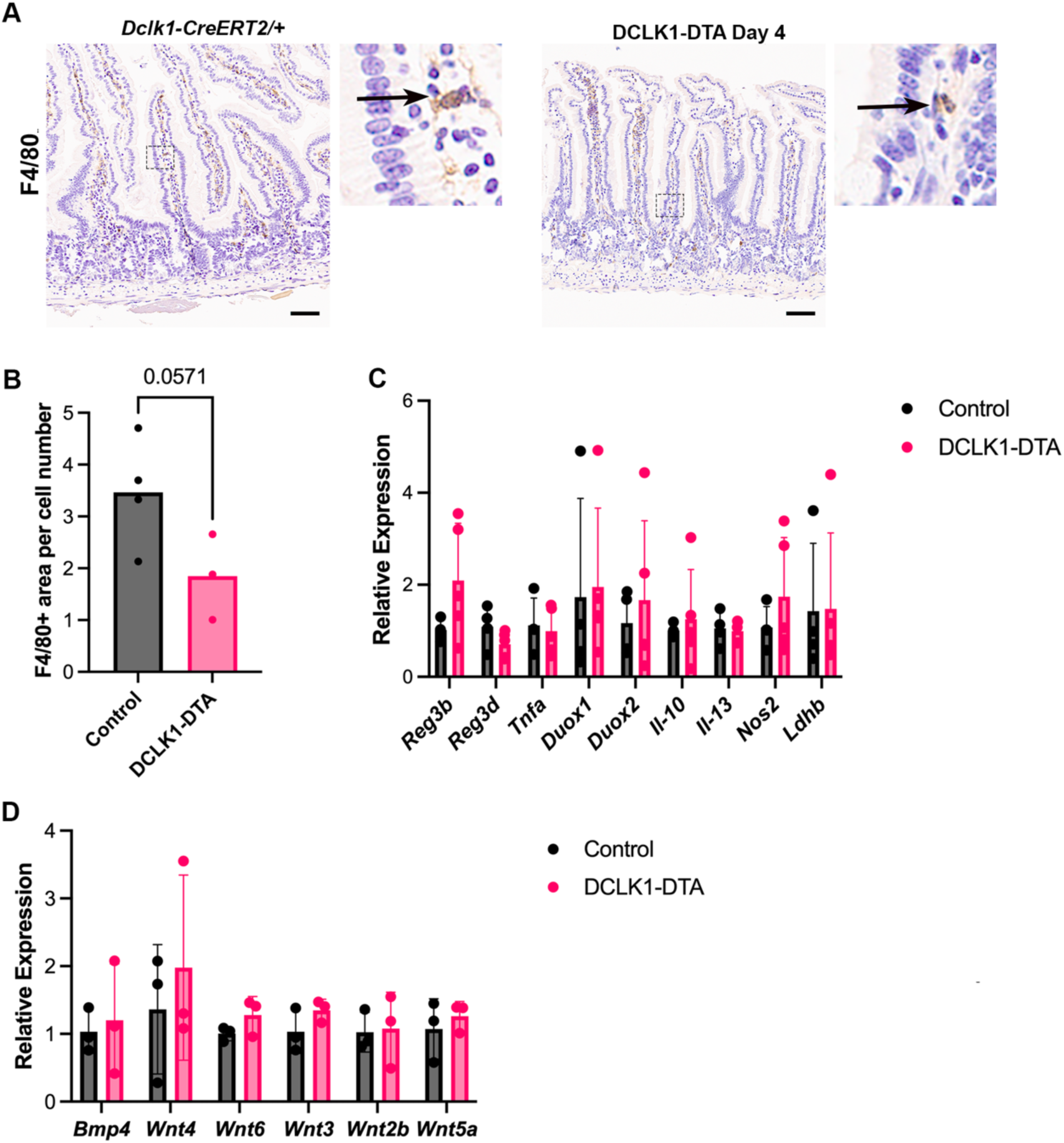
Lack of changes in proinflammatory and growth factor signaling molecules in tuft-cell deleted mice. **(A)** Immunohistochemistry for F4/80, a marker of macrophage and dendritic cells, in duodenum. Scale bars = 50 µm. **(B)** Quantification of F4/80+ cell area per mucosal cell nucleus number. No statistical difference is detected by Mann-Whitney test. **(C)** Transcription levels of proinflammatory markers in scraped mucosa. No difference is detected by two-way ANOVA. **(D)** Transcription levels of secretory growth factors that influence epithelial cell proliferation.

Proinflammatory marker genes were assessed by qRT-PCR using RNA from scraped jejunal mucosa. Transcription levels were assessed for regeneration family members (Reg3-beta and –gamma), cytokines such as tumor necrosis factor-alpha (*Tnfa*), interleukin 10 (*Il-10*), and interleukin 13 (*Il-13*), inflammatory marker enzymes including lactate dehydrogenase B (*ldhb*), *Duox1* and *Duox2*, and inducible nitric oxide synthase (*Nos2*). There was no significant difference between DCLK1-DTA mice on day 4 versus controls, indicating that the intestinal mucosa did not initiate an inflammatory response when tuft cells were acutely deleted (**Figure 7C**). Due to the decreased epithelial proliferation in the DCLK1-DTA mice, we assayed the expression of growth factors, including Wnt family ligands and a Wnt inhibitor, Bmp4. As Wnt is associated with adult tissue regeneration (34), we sought to determine if lower levels of Wnt or higher Bmp4 could be responsible for the shorter small intestine. However, there was no significant difference in expression of these growth factors between DCLK1-DTA mice and control tissues on day 4 post tamoxifen (**Figure 7D**)

### Biliary tuft cells are depleted along with intestinal tuft cells in the DCLK1-DTA mice

Gallbladder epithelium harbors bile acid-sensing tuft cells, which have distinct transcriptional signatures from intestinal tuft cells and may influence digestive function (35). To assess whether biliary tuft cells are also affected in our DCLK1-DTA mice, whole mount gallbladders were immunostained to identify tuft cells. Both DCLK1-DTA mice and control littermates, which carry a heterozygous GFP knocked-in allele (*Dclk1-GFP-CreER^T2^/+*), were immunostained for GFP and DCLK1 (**Figure 8A and 8B**). Additional staining with phalloidin (F-actin) identified dense “tuft” microvilli and actin rootlet structures of biliary tuft cells similar to those of intestinal tuft cells (36). *Dclk1-GFP-CreER^T2^/+* control tissues demonstrated GFP+ / DCLK1+ tuft cells in the gallbladder and intestines, while DCLK1-DTA mouse gallbladders that were harvested 4 days post-tamoxifen treatment showed few tuft cells (**Figure 8A**). These observations indicate that this mouse model effectively diminishes the tuft cell populations in biliary and intestinal epithelia. Ussing chambered murine gallbladders demonstrated forskolin-induced *I*_sc_ increases accompanied with a decrease in *R*_t_, likely representing chloride secretion (Figure 8C). This response was significantly reduced by tuft cell depletion, suggesting that biliary tuft cell loss may influence gallbladder function and decrease nutrient absorption process in the small intestine.

**Figure 8.**
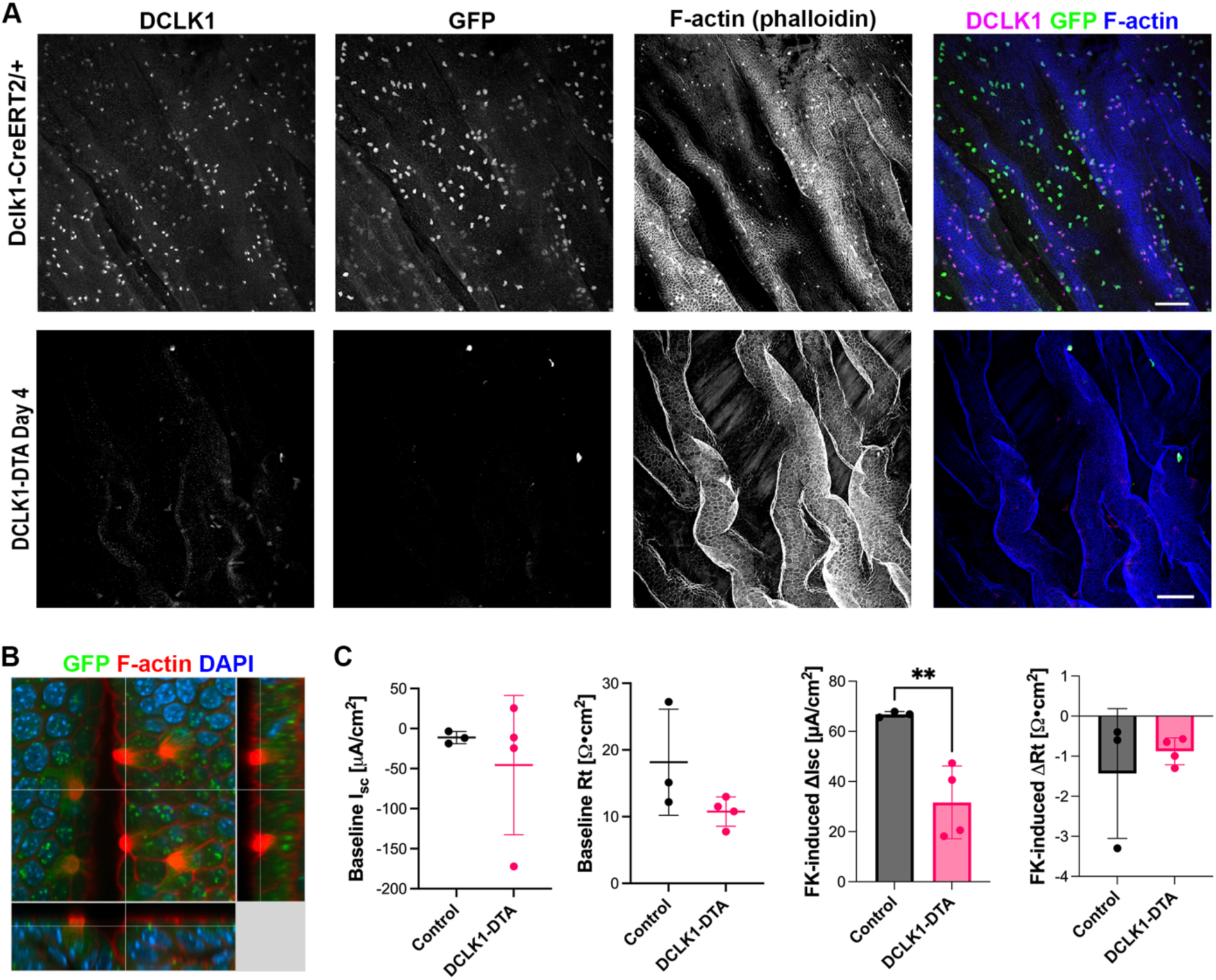
Acute depletion of biliary tuft cells in DCLK1-DTA mice. **(A)** Immunostaining for DCLK1 and GFP in whole-mounted gallbladders 4 days post tamoxifen injection. F-actin and nuclei are stained with phalloidin and Hoechst, respectively. Scale bars = 50 µm. **(B)** Three-dimensional reconstruction of confocal images of tuft cells in control mouse gallbladder. Dense F-actin microvilli and actin rootlet structure is identified in GFP+ tuft cells. **(C)** Ion transport activities of gallbladders in Ussing chamber system.

## DISCUSSION

The tuft cell population is lost in several gastrointestinal diseases and this loss is typically coincident with other deficits within the intestine, including malabsorption. In this study, we sought to highlight the effects of acute tuft cell deletion independent of a chronic disease state utilizing a Diphtheria toxin subunit-A expression system in *Dclk1*-expressing cells (DCLK1-DTA). Following a single tamoxifen injection, tuft cell numbers were decreased in a time-dependent manner. Tuft cells disappeared between 2 and 4 days after injection and recovered to normal levels by day 7. Our data demonstrates that acute tuft cell deletion induces body weight loss due to malabsorption within the murine small intestine. We identified significant changes in immune cell composition, an increase in aberrant secretory cell lineage differentiation, a decrease in proliferative crypt height, and reduced glucose absorption along with small intestinal shortening.

Intestinal shortening is usually observed in colitis with additional hallmarks including inflammation of the colon as well as overall body weight loss (37, 38). In this study, however, we only observed a significant shortening in the length of the small intestine, with no difference in the length of the colon or no pathological signs of inflammation (**Figure 1B–C, 7A–C**). Intestinal shortening following tuft cell deletion could explain the body weight loss due to a reduced absorptive area as compared to controls (**Figure 1A**). The correlation of small intestinal length and tuft cell number has been identified in another mouse model: the deletion of A20, a deubiquitinase, which inhibits the release of IL-13 from ILC2s, leads to constitutive activation of the type-2 immune response and subsequently results in tuft cell hyperplasia (39). Consequentially, in the A20 deficient mice, the tuft cell hyperplasia coincides with small intestine lengthening; the opposite of what was observed in our tuft cell deletion model. In the present study, exogenous recombinant mouse IL-25 (mIL-25) could overcome the phenotype of DTA-induced tuft cell deletion in mice (**Figure 1B**). Supplementation with mIL-25, which is secreted by tuft cells under homeostasis, to DCLK1-DTA mice prevented the shortening of the intestine and body weight loss. This confirms that tuft cells play in extensive role in small intestinal remodeling, a relatively unexplored function of tuft cells. A significant decrease in proliferative cells in the small intestine after acute tuft cell deletion supports the hypothesis that tuft cells are important for maintaining intestinal epithelial proliferation. We hypothesize that compensation occurs in the germline tuft cell deletion model, however, further studies are required to identify the underlying mechanism (**Figure 2A–F**).

Nutrient absorptive function in the small intestine largely depends on the activity of transporters localized to the apical membrane of enterocytes. SGLT1 malfunction is implicated in the absorptive deficit of sodium, glucose, and water in the intestine (40). In our functional assays, DCLK1-DTA mice demonstrated significantly impaired SGLT1 activity after the acute tuft cell deletion without transcriptional changes in *Slc5a1* or *Tas1r2/Tas1r3* (**Figure 3A–C, 3F– H**). Currently, there is little known about the connection between tuft cells and SGLT1 function or glucose metabolism. Tuft cells are positively correlated with improved glucose metabolism in obesity models (41). This function, as well as tuft cells’ involvement in maintaining the mucosal barrier function in the intestine (42), may impact on SGLT1 activity on the apical membrane of enterocytes. The precise mechanism of SGLT1 regulation by the tuft cells remains open for investigation. The loss of biliary tuft cells in the DCLK1-DTA mice in a similar time course to intestinal tuft cells, likely decreased overall nutrition absorption and contributed to body weight loss (**Figure 8**).

Active chloride excretion drives water secretion into the intestinal lumen (43). This helps maintain intestinal barrier function and assists in parasite clearance (7, 8). Epithelial chloride secretion is mostly maintained by two intracellular signaling molecules, cAMP and Ca^2+^, and two membrane transporters, CFTR and CaCCs (44, 45). CFTR serves as the major source of chloride secretion in the murine intestine, and its disruption can lead to inflammation, dysbiosis, and cancers (46). Acutely deleting tuft cells in the intestine led to reduced chloride secretion; in particular, the CFTR-dependent portion of our study (**Figure 3E-F**). This correlates to the reduction of crypt length in tuft cell deleted mice where CFTR is highly expressed (**Figure 2A– B**). Although there was no evidence of inflammation in our acute model, this observation may explain why the intestine is highly susceptible to inflammation upon tuft cell deletion as previously reported with ChAT KO, Pou2f3 KO, and FLARE25 mice (11, 12, 47). In CFTR KO mouse models, goblet cell hyperplasia in the intestinal mucosa is a secondary consequence of the absence of CFTR (46). In the present study TFF3+ cells remained at relatively steady numbers between control and DCLK1-DTA tuft cell deleted mice. Instead, we observed an increase in aberrant goblet/Paneth mixed cell lineage signatures in the small intestine (**Figure 4A-F**).

Goblet and Paneth cells are actively involved in protecting the intestinal barrier from invading pathogens. Goblet cell hyperplasia occurs during the type-2 immune response to aid in the clearing of invading pathogens (12). In a LYZ-1 deficient mouse model, the intestine maintains homeostasis by activating a type-2 immune response, which is dependent on tuft cell expansion in the absence of Paneth cells (48). In our tuft cell depletion model, we observed an increase in lysozyme production in goblet-like cells in villus regions (**Figure 4B**). Mature Paneth cells normally localize to the base of crypts, and mature goblet cells to upper villus regions, which do not typically express lysozyme. Stem cells differentiate into goblet and Paneth cells through Wnt/Notch signaling (49). Recent single-cell RNA-sequencing data have revealed that transcriptional differences between these two secretory cells are minimal and there is likely a common precursor to Paneth and goblet cell lineages (49). The increased frequency of goblet/Paneth mixed type cells in the villi of DCLK1-DTA mice at day 4 led us to suspect that these cells were immature secretory cells, in which terminal differentiation was disrupted by tuft cell loss. This phenomenon has also been seen in Sox-9; Villin-Cre mice, where tamoxifen-induced conditional deletion of Sox9-dependent Paneth cells in the intestine resulted in LYZ+ granules being detected in villus epithelial cells beyond their normal distribution in the crypts (28). This staining pattern phenocopied in *Pou2f3* germline KO mice, where we also observed these mislocalized Paneth cells in the villus (**Figure 4C**). This observation further bolstered our hypothesis that the rise in the goblet/Paneth progenitor cells after tuft cell loss is likely a protective response. This is also supported by the increased frequency of enteroendocrine cells (EECs) in the DCLK1-DTA mouse day 4 intestine (**Figure 5A-D**). EEC is another chemosensory cell lineage and immune response modulator (48). Cholecystokinin (CCK), a hormone secreted by a subset of EECs in response to luminal nutrients, is involved in resolving helminth infection (29). CCK+ cell hyperplasia leads to reduced feeding, therefore reducing fat-produced inflammatory adipokine leptin, which enhance type 2 immune responses to expel parasites from the small intestine. In the present study, specifically CCK+, but not 5-HT+, GIP+, or SOM+, EECs were significantly increased following tuft cell deletion (**Figure 5C and D**). This result highlights the potential for EECs to compensate for the void left by acute deletion of tuft cells.

Tuft cells trigger immune responses principally via IL-25 secretion which activates ILC2s in the lamina propria as a first step towards parasite eradication in the intestine (11). ILC2s, in addition to secreting IL-13, also recruit macrophages, eosinophils, and granulocytes as part of the immune response to invading helminths (50, 51). In the present study, we observed a decrease in the number of mast cells and leukocytes in the intestine but no change in T-cells and macrophages after acute tuft cell deletion (**Figure 6 and 7**). These observations suggest that tuft cells play a role in maintaining some innate immune cell populations within the mucosa even when though there is no parasitic presence in the intestine. Inevitably, this may also play a part in the increased susceptibility of the intestine, after tuft cell loss, to inflammation when challenged (11, 12, 47).

In summary, this study shows that acute tuft cell deletion induces a host of small intestine remodeling effects, highlighting new roles for tuft cells in nutrient absorption. We showed that tuft cell loss led to a decrease in the body weight and a shortening of the small intestine in DCLK1-DTA mice, which was restored by the administration of IL-25. Glucose absorption within the jejunum was hampered, highlighting a novel connection between tuft cells and SGLT1. This mouse model could be used to deepen our understanding of glucose absorption in the intestine through SGLT1 regulation in a tuft-cell-mediated circuit. Furthermore, reduced chloride secretion, decreased eosinophils, and mast cells showed an abridged ability for the intestine to respond to invading pathogens or allergen challenges. However, adaptive responses such as increased enteroendocrine cells as well as the rise of an abnormal secretory lineage displaying goblet and Paneth cell markers may compensate the intestine homeostasis. Broadly, alterations in secretory cell lineages could serve as a focal point for understanding intestinal diseases and may provide insight into the treatment of such diseases.

## Abbreviations

5-HT: 5-hydroxytryptamine
ACTG1: gamma-actin
CaCC: calcium-activated chloride channel
CCh: carbachol
CCK: cholecystokinin
CFTR: cystic fibrosis transmembrane conductance regulator
CHGA: chromogranin-A
DCLK1: doublecortin like kinase 1
DTA: diphtheria toxin fragment A
EpCAM: epithelial cell adhesion molecule
GIP: gastric inhibitory peptide
ILC2: group 2 innate lymphoid cell
IP: intraperitoneal
LYZ: lysozyme
MCPT1: mast cell protease 1
MMP7: matrix metalloprotease 7
MxIF: multiplex immunofluorescence
PAS: periodic acid–Schiff
SGLT1: sodium-dependent glucose transporter 1
SST: somatostatin
TFF3: trefoil factor 3

## Acknowledgements

We thank the Vanderbilt Digital Histology Shared Resource (VA Shared Equipment grant 1IS1BX003097), the Vanderbilt Genome Editing Resource (RRID: SCR_018826) supported by the Cancer Center Support Grant (<u>CA68485</u>), and the Vanderbilt Translational Pathology

Shared Resource supported by NCI/NIH Cancer Center Support Grant P30 CA68485. Core Services performed through VUMC Digestive Disease Research Center (DDRC) supported by NIH grant P30DK058404.

